# First vs recurrent episode symptomatology in Major Depressive Disorder and its relation to brain function and structure: a network approach

**DOI:** 10.1101/2025.04.12.648523

**Authors:** Diego Angeles-Valdez, M.J. van Tol, Eduardo A. Garza-Villarreal

## Abstract

**Background.:** Major Depressive Disorder (MDD) is a prevalent psychiatric disorder. At least half of the patients who recover from a first depressive episode, will experience a relapse. Therefore, understanding the underlying mechanisms supporting relapse is a clinical urgency that could be informed by studying complex brain-behavior associations. Here, we investigated how the relationships between depressive symptomatology and regional brain characteristics differed between people with first depressive episode vs recurrent depression.

**Methods.:** We used REST-meta-MDD data from the DIRECT consortium. We focused on comparing global and local network properties between first (n=239) and recurrent episode (n=179) on: (i) symptom network, (ii) brain structural (VBM) and functional networks (ALFF, ReHO), and (iii) integrated symptoms network and brain characteristics using the psychopathology and multimodal network approach.

**Results.:** Symptom network analysis showed high values of strength centrality for *“Insomnia: Early Hours of the Morning”* and *“General somatic symptoms”* at recurrence compared to the first episode. Also, differences in global strength in the integrated symptom-brain network (measured with ReHo metric) (S=2.09 *p*= 0.042). Finally, we found the edge of specific symptom-brain links, including insomnia and somatic symptoms-, to differ between the first episode and recurrence.

**Conclusions.:** For symptom networks, local but not global properties differentiated first from recurrent episode MDD, with specially stronger relations of insomnia and somatic symptoms in recurrent episode depression. For integrated symptom-brain networks, global strength of the network reflecting regional functional integrity (ReHO) was related to recurrence. This suggests that symptoms have relevance for understanding the complex brain-symptom relations underpinning recurrence of depression.

## 1. Introduction

Major depressive disorder (MDD) constitutes a significant factor contributing to the global burden of disability ^1^, affecting approximately 19% of the global population and is one of the most prevalent mental health concerns. MDD is a heterogeneous disorder ^2^, characterized by large variation in etiology, depressive symptomatology ^3,4^, and its course ^4,5,6^ has been contributed to a large variation in biological findings ^7,8^. The recurrence of MDD is defined as the return of symptoms during a period of recovery ^9,10^, and characterized by persistent suffering, poor general health status, and detrimental effects on psychosocial, educational, occupational, and family functioning ^11^. Moreover, the economic costs associated with depression are significant due to its recurrent nature, an increase in physical health problems that affect quality of life, and an increased risk of suicide ^12^. Of all people who suffer from their first episode, around 50% will relapse in a new episode within two years. For people in their second, third or subsequent episode, this risk is significantly higher and may go up to 70-90% ^13^. Clinical comparisons between first-episode and recurrent depression reveals clinical differences ^14,15^ Understanding these differences may provide insights into the factors that contribute to relapse and, consequently, inform the development of preventive interventions ^16^. In recent years, the psychopathology network approach has emerged as a novel method for modeling and understanding mental disorders ^17,18^. This network approach conceptualizes mental disorders as resulting from a complex interaction between symptoms. Typically, items in psychometric scales are used as nodes, connected by edges indicating a statistical relationship between them. Network models examine how symptoms interact and influence each other. Recent studies have used the network approach to investigate the global and local properties of symptoms in psychiatric disorders such as autism ^19^, posttraumatic stress disorder (PTSD) ^20^, bipolar disorder ^21^, borderline personality disorder (BPD) ^22^. In depression, differences in the global structure of symptom networks were also found between patients with current depression compared to subclinical depression and healthy controls ^23^. Additionally, another study reported that persistent depression was characterized by a more closely connected network (higher number of connections) compared to patients in remission depression ^24^.

Depression has been associated with structural and functional brain abnormalities, but findings have not been wholly consistent. This could be owing to clinical heterogeneity between individuals ^7,25,26^. Multi-modal network methodology allows the integration of brain and behavior ^27^ by explicitly making use of individual variability. It has been used to identify links between mental health disorder and brain ^28–30^. However, to the best of our knowledge, there are few studies using integrative networks in depression. Previous findings have established initial links between individual depressive symptoms and cortical thickness in depression ^29^ and in adolescents with depression ^30^, yet only structural MRI measures have been explored. Investigating the differences in psychopathology and networks covering multiple brain properties between first and recurrent episodes may provide a more complete understanding of the relationship between symptoms and brain characteristics.

In the present study, we continue to explore the approach proposed by Hilland et al. (2020) and aim to investigate the global and local network properties of depressive symptomatology at different clinical stages (first depressive episode and recurrent patients), as well as their relationship with two MRI modalities (brain structure and function). We hypothesize that there will be differences in both global and local properties within the symptom network between the first and recurrent depressive episode. In general, we expect that recurrent MDD to have significantly higher global network values (i.e., are more strongly connected) for symptom, brain and integrated networks. The findings of this study will offer crucial insights into clinical symptoms and neural mechanisms underlying MDD, reflecting about the heterogeneous nature of depression. Additionally, the results may contribute to the identification of symptoms-brain links that may allow us to identify patients at higher risk of disease progression, thus improving treatment prospects.

## 2. Methods

### 2.1 Participants

For our study, we used the REST-meta-MDD project dataset based on the Depression Imaging Research Consortium (DIRECT) initiative ^31^, which consists of patients from 17 hospitals in China. The REST-meta-MDD project includes 25 cross-sectional studies, with MRI-, clinical-, and demographic data from 1,300 patients with major depressive disorder (MDD) and 1,128 non-affected controls (NACs) ^31^. During data collection, the patient’s clinical status was recorded (first episode or recurrence) ^32^, including whether the patient’s previous and current episodes were diagnosed as MDD according to ICD-10 or DSM-IV ^33,34^. For the present analysis, we selected a subsample based on participants who had MRI data and clinical measures. Our final sample was 418 MDD patients (First episode = 239 [162 female] and Recurrence = 179 [110 female]). This study was approved by the Consortium’s Institutional Review Board (IRB), approval number H19045, of the Institute of Psychology, Chinese Academy of Sciences. All participants provided written informed consent. A flowchart detailing the sample selection process for the study design is available in Figure 1a.

**Figure 1.**
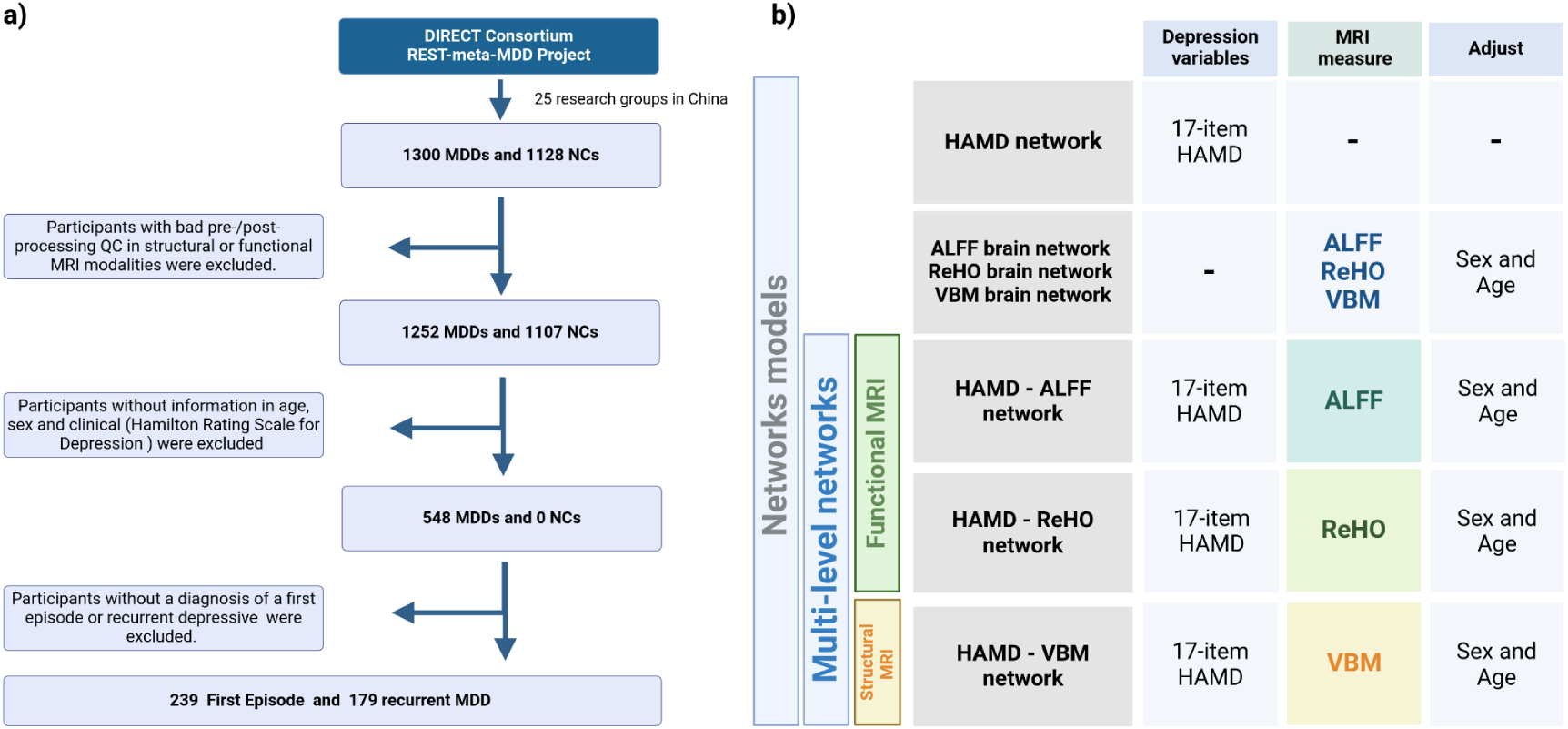
Flow chart of the study population and network models characteristics. **Note: a)** Flow chart of the selection of the study population for network analysis. **b)** Description of symptom network and multi-modal network. HAMD: Hamilton Rating Scale for Depression; ALFF: Amplitude of Low Frequency Fluctuations; ReHo: Regional Homogeneity; VBM: Voxel-based morphometry. Created with bioRender.

### 2.2 MRI preprocessing

All MRI scans were acquired at each of the hospitals, details on each of the MRI scanners and protocols are in the supplementary material. MRI preprocessing was carried out by the REST-meta-MDD Project of the DIRECT consortium using the following tools. For structural data we used the T1-weighted standard pipeline of the DPARSF software that was applied for image preprocessing ^35^. T1-weighted images were segmented into cerebrospinal fluid (CSF), white matter (WM), and gray matter (GM). The Diffeomorphic Anatomical Registration Through Exponential Lie (DARTEL) tool was then used to transform individual native space into MNI space, and then smoothed by using an 8-mm full-width half-maximum Gaussian kernel. For Voxel-based morphometry analysis (VBM), the VBM 8 toolbox implemented in SPM 8 software was used ^36^. Functional data were preprocessed using a standardized preprocessing pipeline from Data Processing Assistant for Resting-State fMRI (DPARSF) ^37^, and Amplitude of Low Frequency Fluctuations (ALFF) and Regional Homogeneity (ReHo) were estimated for each participant, details of MRI preprocessing can be found in the supplementary material. For this work we used the final processed data for each modality: individual VBM, ALFF and ReHo maps. Finally, to control for multisite MRI variations, we applied a harmonization method with NeuroCombat tool (https://github.com/Jfortin1/ComBatHarmonization) on the brain images ^38^ and we adjusted for variables such as sex, age, and intracranial volume only for the structural images.

### 2.3 Statistical analysis

To study differences in the local and global properties between the first episode and recurrent depression, we used a psychopathological network approach. First, we defined the clinical and brain nodes, then estimated separate clinical and brain network models, and finally combined symptoms and brain in a multi-modal network approach (integrated network) ^27^.

#### 2.3.1 Node selection

For the selection of symptom *nodes* we used the Hamilton Rating Scale for Depression (HAMD) with 17 items describing vegetative/somatic and cognitive symptoms ^39^. For the selection of brain nodes, we extracted two functional MRI metrics: Amplitude of low-frequency fluctuations (ALFF), defined as the total power for each voxel of the brain, and Regional homogeneity (ReHo), defined as the correlation of the time series of a voxel with a local neighboring voxel ^40^, and for structural metric we extracted brain volume (gray matter) using Voxel-Based Morphometry (VBM). We used six Regions of Interest (ROIs) related to MDD, based on previous studies ^29,41,42^ (medial Prefrontal cortex, Cingulate cortex, Insula, Fusiform, and Hippocampus), and in order to explore the integrated network relationship the thalamus values ^43,44^ was also extracted. We created MRI ROIs using the WFU PickAtlas toolbox (http://fmri.wfubmc.edu/software/PickAtlas).

#### 2.3.2 Network estimation

We estimated seven network models for each group separately: 1) a symptom network (HAMD network), 2) brain networks based on three MRI measures: ALFF, ReHO, and brain volume, and 3) an integrated network to combine brain and behavior which individual depression symptoms ^27^. The estimation of the models is described in the next section (see Figure 1b for all network models).

##### 2.3.2.1 Clinical and brain network

First, we estimate a separate regularized partial correlation network for clinical symptoms and brain nodes. For the symptom network, we included 17 symptom nodes, each representing an item from the HAMD scale, and we used the polychoric correlation matrix to consider the ordinal nature of the data, and for the brain network, we used six brain regions as nodes. Four different network structures (Clinical and VBM, ALFF and ReHO metrics) were generated by independent estimation for each group (First episode and recurrence), where the edges between nodes represent conditional independence relationships between nodes. The network models were regularized with LASSO (least absolute shrinkage and selection operator) ^45^ was used which has high specificity. Finally, the best-fit network models with the lowest EBIC value were selected with hyperparameter gamma γ set to 0.5 and the min.lambda was set to 0.1 ^46,47^.

##### 2.3.2.2 Integrated Network

Finally, an integrated network approach was used to connect the brain and individual depression symptoms. We estimated an integrated network with 23 nodes representing the six *brain nodes* and the 17 items of the HAMD scale. The network structures were generated through a separate estimation for each group (First episode and recurrence), one for each MRI measure (brain volume, *gray + white matter*), ALFF and ReHO); here, the brain variables were modeled as a Gaussian distribution. For comparability of results, the networks were estimated using the same regularization parameters as those described in the previous section.To calculate the brain correlation matrices, we used z-residuals adjusted for the effects of sex, age, and intracranial volume on the structural measures^42,48^. Then, we estimated the regularized partial correlation network. All networks were estimated using the *bootnet* in R package ^45^.

### 2.4 Network comparison

In order to compare global and local properties between groups (first episode and recurrence). First, we performed a *Network Comparison Test (NCT)* using a permutation test with 4000 iterations ^49^ to compare the global strength and network structure between first episode and recurrence. A statistical significance was set at 5%. Local properties comparisons are described in the next section.

### 2.5 Metrics of network centrality

In psychopathological networks, centrality measures are commonly calculated to determine the relevance of nodes within the network structure. We computed and compared the most frequently used centrality measures within the network structure, only for the “unimodal” clinical and brain networks: (i) *strength* is a measure of how strongly a node is directly connected to other nodes, (ii) *closeness* examines how strongly a node is indirectly connected to other nodes; and (iii) *betweenness* examines the importance of a node in connecting other nodes ^45,50^. Finally, to determine which symptoms were most different between first episode and recurrence, differences in invariance of centrality measures were calculated using NCT with 4000 iterations.

### 2.6 Network stability

After estimating networks, a bootstrapping method was applied to check the accuracy and stability of the edges strengths and centrality measures. First, we estimated confidence intervals (CIs) with 4000 interactions to determine whether the edge weights were not zero ^45^. Then, to investigate the stability of the centrality measures, we used the case-dropping bootstrap to estimate the correlation between the original centrality coefficients and those obtained from different subsets (75%, 50%, and 25%), the estimated centrality measures were considered ideally stable if the correlation stability coefficient *(CS coefficient*) remained at 0.25 ^45^.

## 3. Results

### 3.1 Demographic and clinical measures

The final sample included 418 participants with MDD with first-episode (n=239, female = 162), and recurrence MDD (n=179, female = 110). Illness duration from onset was shorter in the first-episode group (Mean = 9.62 months, SD = 13.5) compared to the recurrence group (Mean = 85.5 months, SD = 86.1; t(183.99) = −11.65, *p*<0.001*). Additionally, the first-episode group showed a higher mean Hamilton scale score (22.6, SD = 6.05) than the recurrence group (18.7, SD = 8.14; t(315.29) = 5.42, *p*<0.001*), indicating more severe depressive symptoms at the time of assessment in the first episode group. Descriptive characteristics of the sample are provided in *Table 1*.

**Table 1.**
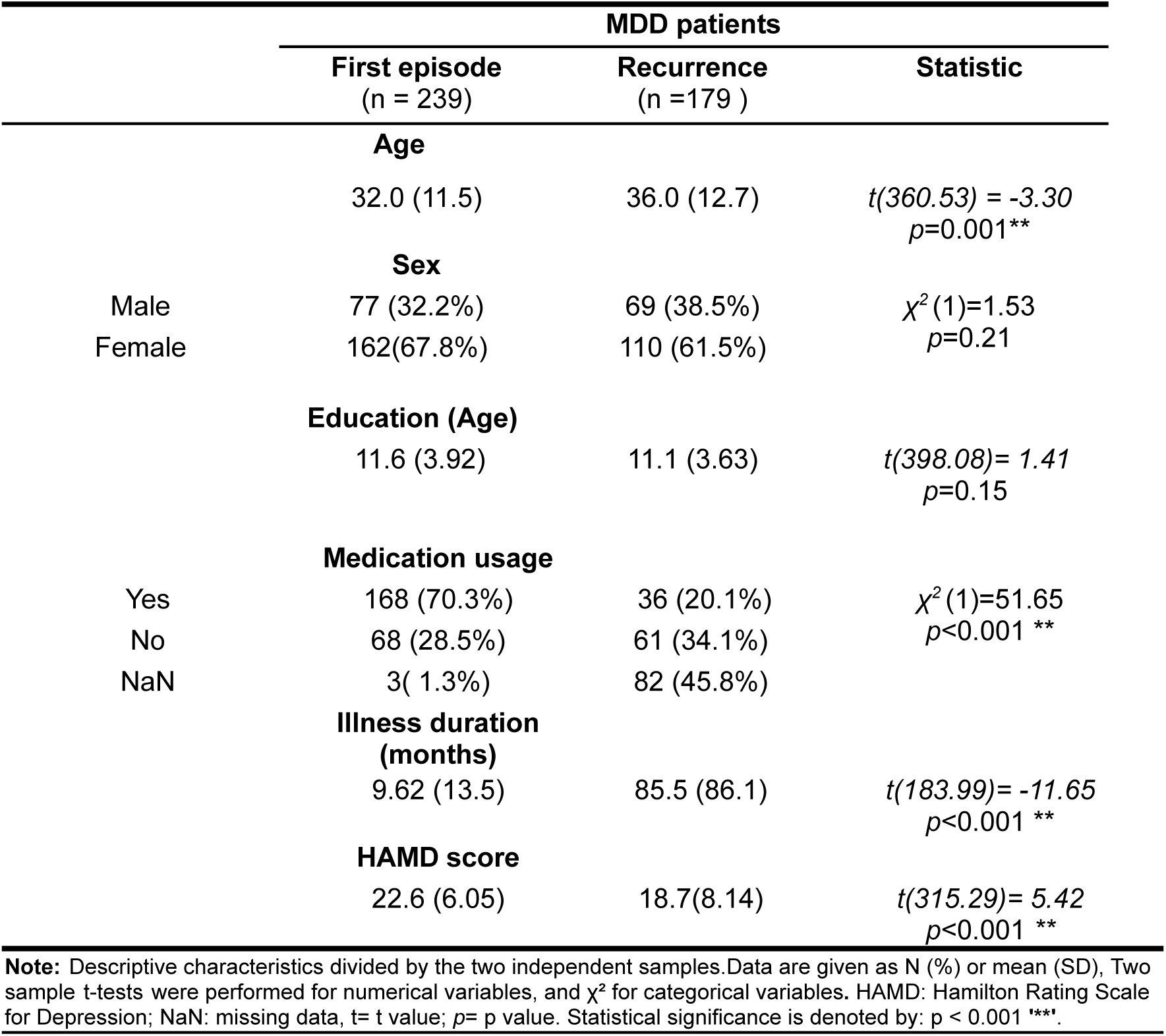
Descriptives and clinical characteristics of the total sample.

#### 3.1.1 Clinical network

For the symptom network, the networks of the first and recurrent episode groups were both mostly positively connected (Figure 2). The number of non-zero edges was higher for the recurrence group compared to the first episode group (70/136, 33/136 edges), this means the recurrence group is more connected. However, the recurrence network seems to be more connected on visual inspection, the network structure test showed no significant differences were observed (M=0.27, *p*=0.15). Furthermore, no significant differences between the first episode and recurrent depression were observed for global strength (S=3.12, *p*=0.11).

**Figure 2.**
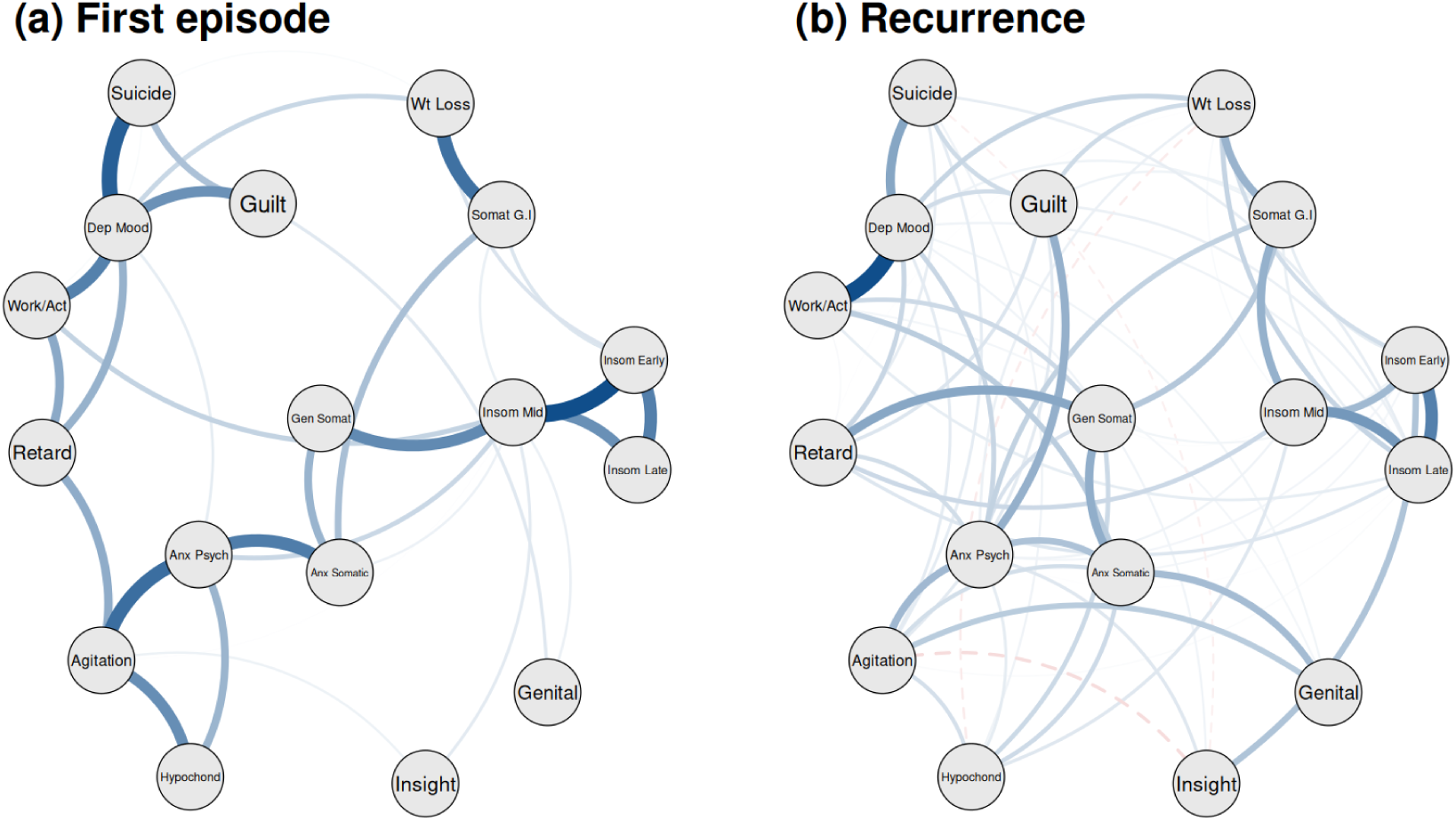
Network structure of the Hamilton Rating Scale for Depression. **Note** Network structure of depressive symptomatology **(a)** first episode, and **(b)** Recurrence. The nodes correspond to the item in the Hamilton Depression Rating Scale (HAMD). Positive and negative connections are represented by blue solid and red dotted edges. Dep Mood: Depressed Mood; Guilt: Feelings of guilt; Suicide: Suicide; Insom Early: Insomnia: early in the night; Insom Mid: Insomnia: middle of the night; Insom Late: Insomnia: early hours of the morning; Work/Act: Work and activities; Retard: Retardation; Agitation: Agitation; Anx Psych: Anxiety psychic; Anx Somatic: Anxiety somatic; Somat G.I: Somatic symptoms gastro-intestinal; Gen Somat: General Somatic symptoms; Genital: Genital symptoms; Hypochond: Hypochondriasis; Wt Loss: Loss of weight.

#### 3.1.2 Brain network

The brain networks were estimated for each of the MRI measures (ALFF, ReHO, and VBM). The majority of edges are consistently preserved across all brain models, particularly in the functional models (ALFF and ReHO brain networks). We did not observe significant differences in global strength and network structure for any of the MRI modalities. The global characteristics and network comparison test of the symptom and brain networks are presented in the supplementary material.

#### 3.1.3 Integrated network

In order to examine the relationship between depressive symptomatology and three MRI measures in individuals with first-episode and recurrent episodes, we have combined these variables into integrated networks (clinical + MRI measure). Figure 4 shows the integrated networks. When we integrated clinical symptoms with brain measures in the first episode and recurrence, positive connections were predominantly maintained for all integrated networks. No difference in network structure test was found for the HAMD-ALFF network (M=0.25, *p*= 0.29), the HAMD-ReHO network (M=0.18, *p*= 0.36) and the HAMD-VBM network (M=0.23, *p*=0.38). However, when we compared the differences in global strength, we only found significant differences in the HAMD-ReHO network (S=2.09, *p*= 0.042), where people with recurrent depression showed higher global strength. The general characteristics of the integrative networks are listed in supplementary material. This result indicates that in our sample, patients with recurrent depression have higher global strength compared with the first episode group in an integrated symptom-brain network where MRI functional measures are represented.

### 3.2 Differences in local properties

#### 3.2.1 Clinical network

We compared strength, closeness, and betweenness as local properties to identify the most relevant nodes within the network between the first episode and recurrent MDD groups. Figure 3 *shows the local properties for the HAMD network model,* specifying the standardized node items of the Hamilton scale. In the symptom network, the Network Comparison Test (NCT) indicated higher local strength in the recurrence group, particularly for Item 6, *“Insomnia: Early Hours of the Morning’’* (*p*=0.011) and Item 13, *“General Somatic Symptom”* (*p*=0.049), these symptoms were more important nodes in the recurrence group, (more details of centrality test in supplementary material). No between group differences were observed for closeness and betweenness.

**Figure 3.**
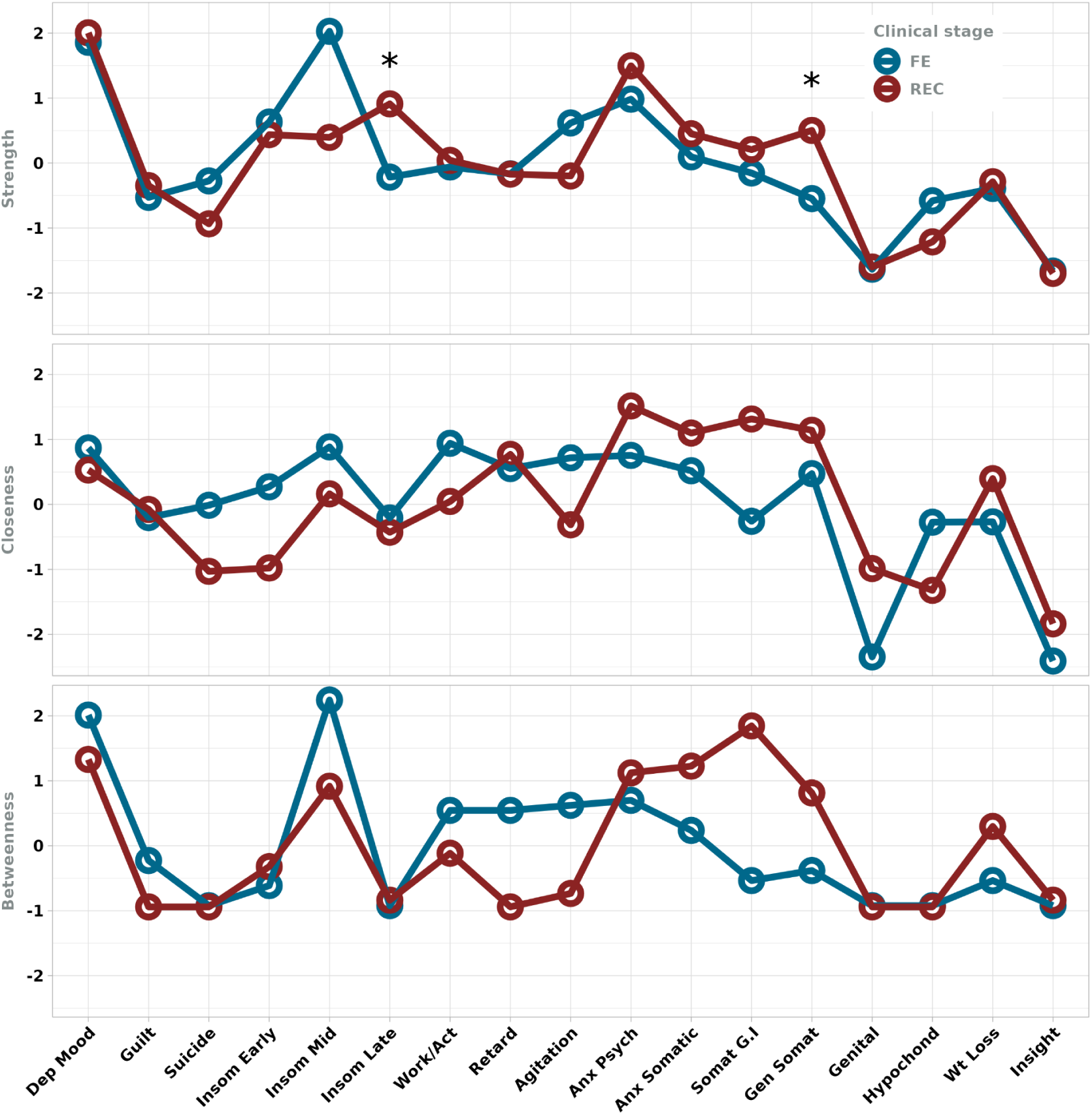
Centrality metrics of clinical depression network. **Note:** Three standardized centrality metrics (Strength, Closeness, and Betweenness) for HAMD network model. Statistical significance is denoted by: p < 0.05 ‘*’. FE:First episode; REC:Recurrence; Dep Mood: Depressed Mood; Guilt: Feelings of guilt; Suicide: Suicide; Insom Early: Insomnia: early in the night; Insom Mid: Insomnia: middle of the night; Insom Late: Insomnia: early hours of the morning; Work/Act: Work and activities; Retard: Retardation; Agitation: Agitation; Anx Psych: Anxiety psychic; Anx Somatic: Anxiety somatic; Somat G.I: Somatic symptoms gastro-intestinal; Gen Somat: General Somatic symptoms; Genital: Genital symptoms; Hypochond: Hypochondriasis; Wt Loss: Loss of weight.

#### 3.2.2 Brain networks

For the brain networks for all modalities, we did not observe statistical differences in local properties; more details in the supplementary material for all comparisons.

#### 3.2.3 Differences in the depression symptoms-brain link

After estimating the integrated network, we investigated whether links between clinical symptoms and MRI measures differ between the first episode (reference group) and recurrence. A permutation test based on resampling was used to compare the strength of edge links between the first episode versus recurrence networks of the symptom-brain networks ^49^. Based on our clinical findings we focus on two symptoms including “*Insomnia: Early morning hours*” and “*General somatic symptoms*,” and their associations with specific brain regions. Two significant differences were found between the groups: (i) between *“Insomnia: Early morning hours - mOFC”* in the VBM measure (E=0.129, *padj* =0.008), and (ii) *“General somatic symptoms - thalamus*” on the ALFF functional measure (E=0.016, *padj*=0.021), as detailed in Table 2.

**Table 2.**
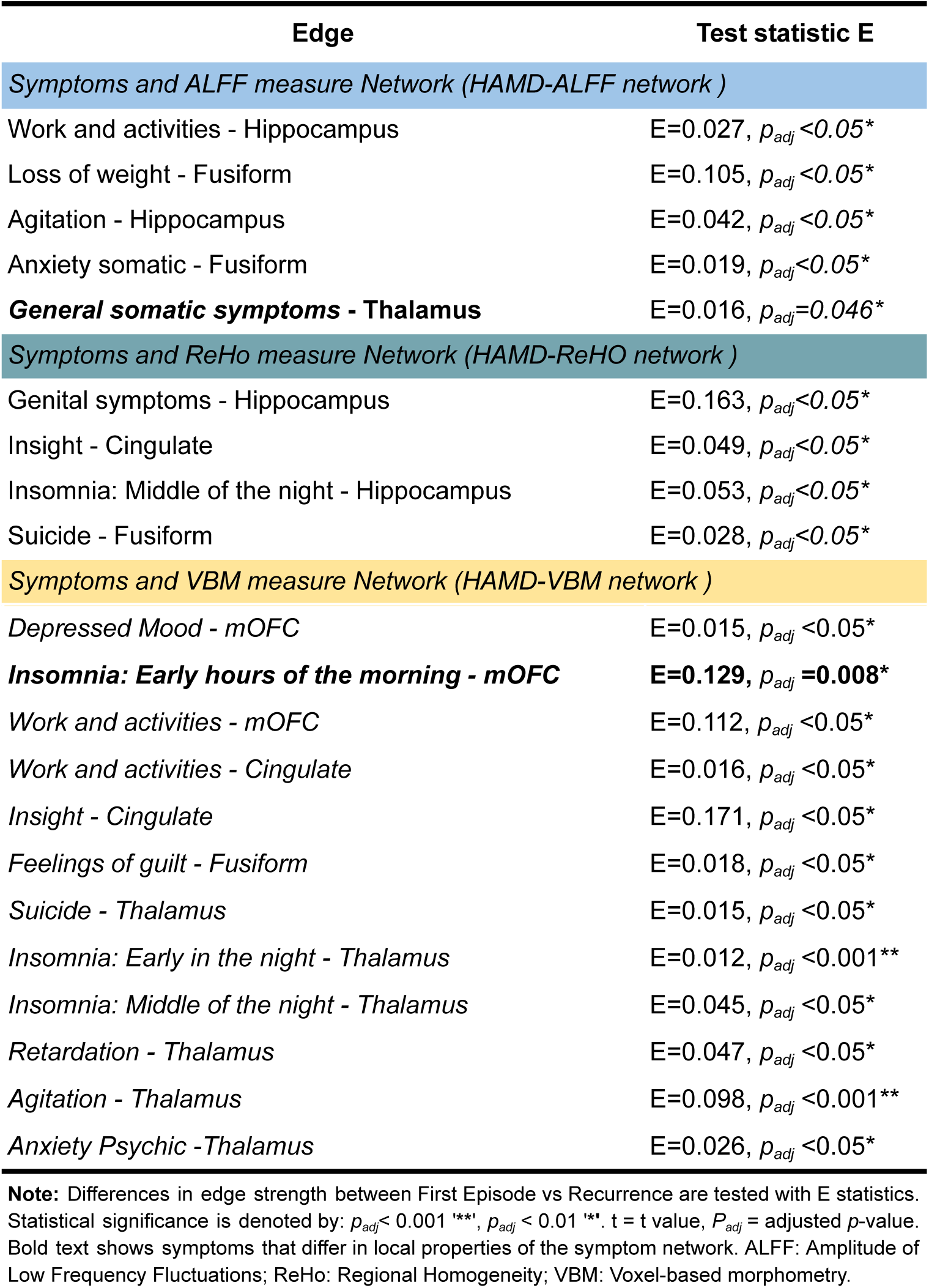
Edge strength comparison of integrated networks.

In addition, the thalamus and hippocampus were the brain nodes with more significant associations (edges) with depressive symptoms. The thalamus was related with *suicide*, *insomnia, retardation, agitation, anxiety psychic and general somatic symptoms (items 3, 4, 5, 8, 9, 10, 13)*. The hippocampus was linked to *insomnia, work and activity*, *agitation*, and *genital symptoms (items 5, 7, 8, 9, 14)*. Here we show that links between clinical symptoms and MRI measures differ between the first episode and recurrence depression, even between MRI measures, where structural measures were related to sleep problems and ALFF measure to somatic symptoms.

### 3.3 Stabilities of strength and centralities

For all estimated network models, we evaluated the stability of the edge strengths with a bootstrapping method. The estimated strengths and bootstrapped means were similar performance for each edge, all bootstrap CIs were greater than zero, this is an advantage of the LASSO regularization we perform. Correlation-Stability Coefficient (CS-Coefficient) indicates whether the centrality metrics remain consistent after the network has been re-estimated with a smaller number of cases. In our estimated networks, the CS-coefficients had a value above the higher cut-off value previously reported (>0.25) ^45^, for strength metrics. However, in some network models, the values of the CS coefficients were close to the cut-off value, thus the findings on centralities should be considered with caution (see the stability performance of the models in supplementary material). Finally, we extracted all edge weights of the integrated networks for each MRI measure, details of which can be found in the supplementary material.

## 4. Discussion

Major depression presents differently between first episodes and recurrences, yet the underlying clinical and biological mechanisms remain poorly understood. In our current study, our goals were to study the network structure of depressive symptoms and brain characteristics of regions previously consistently associated with depression and compare those between patients with first episode vs. recurrent MDD. Using psychopathology and integrated networks derived from a large publicly available dataset, we studied the complex network structure of symptoms and brain characteristics. We expected that clinical and brain networks would show greater connectivity in the recurrent MDD patients. Instead, although we did not find differences in global properties of clinical and brain networks by themselves, we found a more strongly connected network in recurrent MDD when combining symptoms and brain characteristics. These findings suggest that clinical and brain measures are connected in MDD and have relevance for its clinical course. Recurrent MDD may create stable pathological symptom-brain relationships that could contribute to the illness’s chronicity. Furthermore, we also found that insomnia and somatic symptoms seem to be central to recurrent MDD, suggesting these symptoms could serve as possible clinical targets.

### 4.1 Clinical and brain networks

In the symptom network (HAMD network model), no differences in global properties were found between the first episode and recurrence, despite the observation that the symptom network of the relapse group showed a high network density (number of connections). This suggests that the recurrence of depression is unlikely to be attributable to a stronger covariance of symptoms. Thus, the tendency for multiple depressive symptoms to occur simultaneously in patients with a first episode of depression compared to those with recurrent episodes may be linked to similar underlying mechanisms. Previous research found differences in global properties of depression when comparing individuals with current depression to healthy controls ^23^ and when comparing persistent depression with remission condition ^24^. In terms of local network properties, the depressive symptoms identified seem to play different roles between the first episode and recurrence network, specifically *“Insomnia: Early hours of the morning*” and, “*General somatic symptom*” which were more strongly connected in the recurrence network (see Figure 3). Furthermore, a previous study found that insomnia and fatigue or loss symptoms had higher values of the centrality measure (strength) when comparing persistent vs. remitted MDD ^24^. Our findings are consistent with previous literature, suggesting that these symptoms more strongly influence other symptoms and thereby may have a significant impact on the course of depression. Symptom disturbances may occur before a new depressive episode, so it has been suggested that they may be a prodrome, a residual symptom ^51,52^ or is a vulnerability factor for recurrence ^53^. Our results also showed that insomnia symptoms were highly central symptoms in the recurrence group compared with the first episode, which may suggest that they are a risk factor for the development of a new depressive episode ^54,55^. To date, a number of studies have indicated that sleep problems are present in patients with MDD and a recent systematic review and meta-analysis reported that insomnia most significantly predicts the onset of depression (10 studies, OR 2.83, CI 1.55-5.17)^56^. Insomnia has been suggested as a primary risk factor for developing MDD, ^57^ and in our work, we suggest that “*Insomnia: Early hours of the morning*”, which is strongly related to other symptoms, represents a crucial node in recurrence of depression. Consequently, treating insomnia in patients with major depression may be an important treatment target to influence the strong interconnection with other symptoms and to prevent a future depressive episode ^18^. On the other hand, patients with depressive disorders experience a variety of somatic symptoms associated with poor quality of life. It has been observed that patients who experienced a relapse exhibited a higher prevalence of general somatic symptoms compared to those presenting with the initial episode ^14^. Hence these findings showed that somatic symptoms are more closely related to other symptoms when in recurrence thereby treating these symptoms could decrease the risk for relapse.

Regarding the brain networks, we did not find any differences in the global and local properties between the groups (first episode and recurrence) for any of the three MRI modalities (i.e., VBM, ALFF, REHO). When assessing both network structures separately: clinical symptoms and brain information, this provided little information for understanding differences in the course of depression (first episode and recurrence states). Nevertheless, when estimating an integrated network structure, differences between groups were observed. Therefore, we emphasize the need to address depressive symptoms and their relationship with biological levels in order to better understand their impact on the course of depression ^58,59^.

### 4.2 Overall findings in integrated networks

Integrating clinical and brain features provides a more comprehensive understanding of the course of depression. This approach helps us to theorise about the interaction between brain physiology and symptomatology, and shows that symptom heterogeneity is important to consider when trying to understand the brain network properties associated with first vs. recurrent depressive episodes. Symptoms are needed to explain any brain-related variance.

We found global differences only in the functional measure (REHO) we studied, and not in the structural (VBM) measure. The differences observed in the global strength of the HAMD-ReHO network indicate that symptom and local integrity of brain regions commonly associated with depression are more densely or strongly connected in the recurrence group than in the first-episode MDD group. In other words, the strength of the connections between the nodes (clinical and brain regions) serves as an indicator of recurrence of depressive episodes.

Although both integrative networks (HAMD-ALFF and HAMD-ReHO networks) are based on the BOLD signal in each voxel, we only found global differences in the clinical symptoms with ReHO measure (HAMD-ReHO network). This could be attributed to the inherent properties of the used metrics. Specifically, the AlFF value quantifies the amplitude of the BOLD signal representing local brain activity intensity, while ReHo measures capture the local homogeneity of the local neighboring voxels. A notable advantage of ReHo is its capacity to detect unpredicted hemodynamic responses compared to task-based models ^60^. Additionally, assessing local homogeneity with ReHo showed more sensitivity to the identification of brain characteristics related to clinical symptoms. Although ReHo is sensitive to spatial smoothing ^61^, employing a standard pipeline ensures methodological consistency across studies. The differences observed in this study, together with findings from other research, suggest that ReHo holds potential as a marker for depression ^62,63^ and other psychiatric disorders ^62,63^.

Another interesting finding was the differences in the edge strength of the integrated networks for symptoms that differed across the clinical networks, including “*Insomnia: Early morning hours - mOFC*” in HAMD-VBM network and “*General Somatic Symptoms - Thalamus*” in HAMD-ALFF network. These differences were seen after controlling for age and sex between first-episode and recurrence MDD, as shown in Figure 4 and Table 2. Previous studies have explored the brain structural correlates of insomnia symptoms in MDD ^64,65^. The ENIGMA MDD results showed a significant association between more severe insomnia in patients with major depressive disorder (MDD) and structural features in several brain regions, including the medial orbitofrontal cortex ^64^. In our study, we showed structural link between insomnia and mOFC volume in the context of an integrated network analysis. The findings of a stronger association between mOFC and insomnia, in the context of increased global strength of the integrated network in recurrent depression, suggest that this relationship may be relevant to understanding recurrent depression. Future longitudinal studies should investigate whether this is a result of recurrent depression or represents a premorbid vulnerability.

**Figure 4.**
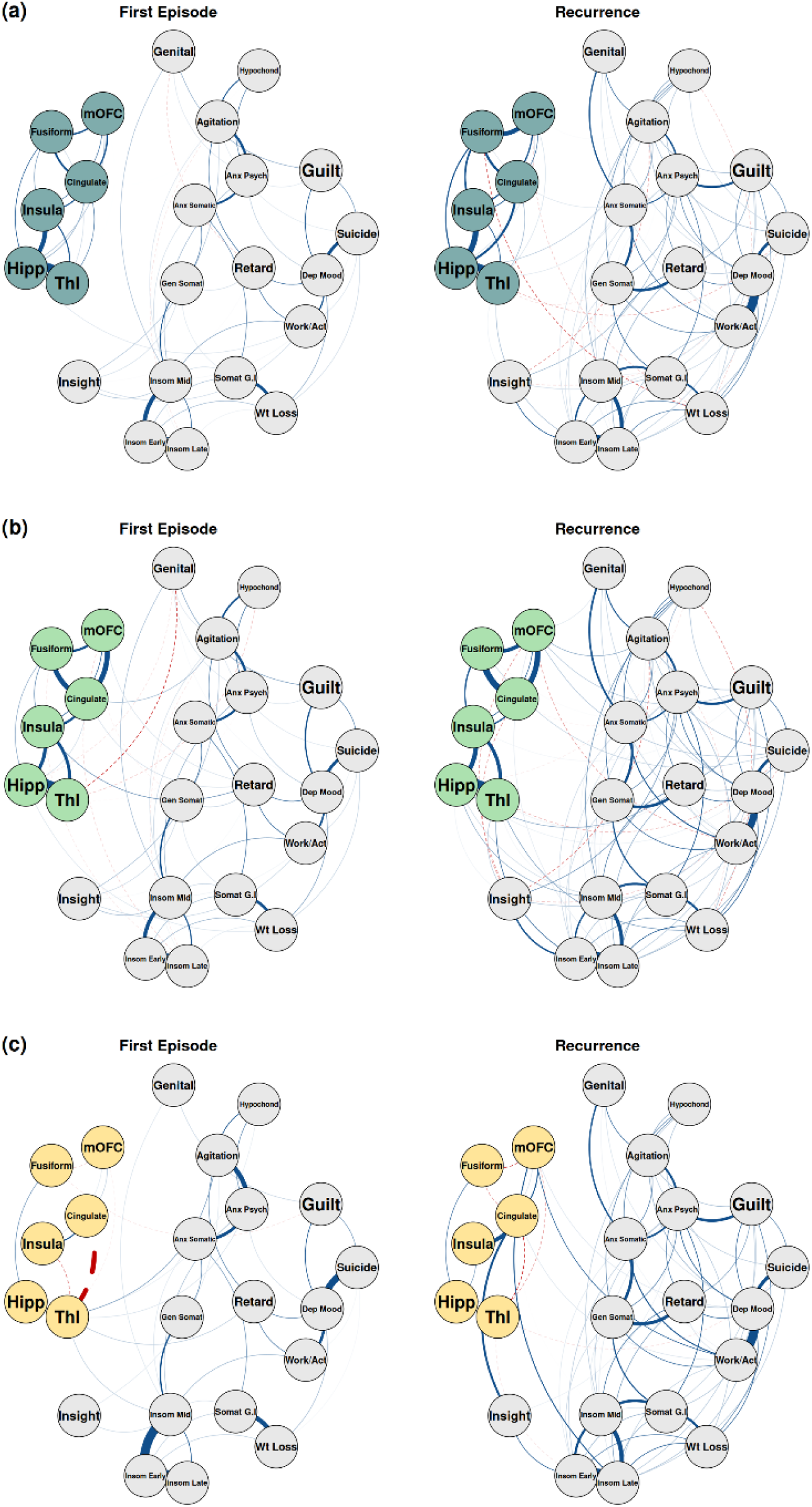
Network structure of Integrated networks. **Note:** The MRI modality are shown in **A)** Amplitude of Low Frequency Fluctuations (blue nodes) and **B)** Regional Homogeneity (green nodes) and structural modality in **C)** Voxel Based Morphometry (yellow nodes). The blue lines represent positive associations and the red dotted lines represent negative associations. Dep Mood: Depressed Mood; Guilt: Feelings of guilt; Suicide: Suicide; Insom Early: Insomnia: early in the night; Insom Mid: Insomnia: middle of the night; Insom Late: Insomnia: early hours of the morning; Work/Act: Work and activities; Retard: Retardation; Agitation Agitation; Anx Psych: Anxiety psychic; Anx Somatic: Anxiety somatic; Somat G.I: Somatic symptoms gastro-intestinal; Gen Somat: General Somatic symptoms; Genital: Genital symptoms; Hypochond: Hypochondriasis; Wt Loss: Loss of weight. mOFC: medial Orbitofrontal Cortex; Hipp: Hippocampus; Thl: Thalamus.

Likewise, although the relationship between “*General somatic symptoms - Thalamus*” has been studied in MDD patients with and without somatic symptoms, where they observed that MDD with somatic symptoms present more severe depressive symptoms associated with Thalamus ^66^, it is also important to note that research on the course of depression, particularly the distinction between first episodes and recurrence, remains poorly studied. Lastly, it is important to emphasise that such associations are speculative in nature, as the estimated associations are undirected and cross-sectional. In addition, integrative networks seem to be a useful approach to effectively identify differences in global properties in depression course (first episode vs recurrence). However, to date, no previous study has examined whether the structure of the integrated network (brain-symptoms) is actually associated with longitudinal prognosis and whether it holds with edges in the brain. This suggests that depression involves biological and clinical measures that are closely linked and that allow us to identify the differences between a first episode and a recurrence.

## 5. Limitations

Our study has several points to consider: while the study of psychopathological networks is a current and novel method, the integration of clinical data with biological measures has been sparsely used so far. A first limitation of the study is that our findings are derived from a cross-sectional study, preventing us from knowing the natural causality of the associations and the role of certain nodes in the relapsing course of depression. Secondly, the categorization of first episode and recurrence was determined by different nosological systems, which may have introduced biases in the comparison, affecting the interpretation and limiting the generalizability of the results. Third, we are familiar with the research on the controversies surrounding the use of centrality measures in psychopathology networks ^67^. However, there is also empirical evidence that centrality indices have been used to describe the importance of symptoms within the network structure^68^. Finally, the estimation of the networks could have been affected by the regularization method and the diversity of clinical characteristics, including medical history and resistance to treatment, or may be influenced in part by cultural and historical contexts, as well as external environmental factors. For these reasons, our results should be considered with caution, and we suggest that future studies use longitudinal designs to assess not only depressive symptomatology but also its relationship with the brain.

## 6. Conclusion

In conclusion, this is the first study to compare first and recurrent episodes of depression using a psychopathology and multimodal network approach, and reveals important differences in depressive symptomatology and brain-behavior associations. Our findings indicate local properties in the symptom network model that distinguish between a first episode and a recurrence of depression. Furthermore, the integrated network analysis allows us to identify specific links (symptoms-brain) that may inform the depression course and suggest that symptom heterogeneity should be taken into account when investigating brain-network properties relevant for the depressive course. Modeling individual depression symptoms with neurobiological measures could be fundamental to unraveling the complex nature of processes in depression.

## Acknowledgments

We thank Prof Mallar Chakravarty (Director of the Computational Neuroanatomy Laboratory, Douglas Research Centre, Montreal, Canada) who provided access to computational tools from his group on the Niagara Compute Cluster from Compute Canada. We also thank the Programa de Maestría y Doctorado en Psicología, Universidad Nacional Autónoma de México (UNAM), for their support and for providing fellowship No. 1003596 from Secretaría de Ciencia, Humanidades, Tecnología e Innovación (SECIHTI). Eduardo A. Garza-Villarreal is funded by PAPIIT-UNAM fund IA201622 and IN213924. We thank Chao-Gan for sharing the REST-meta-MDD Project data from the DIRECT Consortium and Eiko Fried, Said Jimenez and Rozemarijn van Kleef for their comments and support for this paper. Marie-José van Tol is funded through a VIDI-grant (nr. 09150172110019) awarded by the Netherlands Research Council.

